# Neuronal Population Effects of Ketamine on Human Brain Organoids

**DOI:** 10.64898/2026.03.09.710454

**Authors:** Arina A Nikitina, Christian Bustamante Toro, Raymond Gifford, Carolina M Camargo, Barbara Mejía-Cupajita, Kenneth S. Kosik

**Affiliations:** Neuroscience Research Institute and Department of Molecular, Cellular and Developmental Biology, University of California, Santa Barbara, Santa Barbara, CA 93106, USA; Department of Neurobiology, Weizmann Institute of Science, Rehovot, 76100, Israel; Amgen Inc., Neuroscience Research, Thousand Oaks, CA, USA

## Abstract

Ketamine’s rapid neuropsychiatric actions emerge from interactions that span receptors, cells, and circuits, but their net effects on human neuronal population dynamics remain incompletely defined. Here we combine human dorsal forebrain organoids with high-density microelectrode arrays (MEAs) to quantify ketamine’s effects from spikes to networks. In 6-month-old organoids, acute ketamine (20□μg/mL) abolished population bursting while neuronal firing continued mostly unchanged. Spike sorting revealed that mean firing rates declined but not silenced after ketamine Reductions were concentrated within a subset of burst-driver units previously defined as “backbone”. Functional connectivity, estimated with the spike time tiling coefficient (STTC), decreased globally after ketamine. Backbone units displayed elevated connectivity at baseline but were functionally disconnected by ketamine. Graph construction from STTC uncovered widespread network reconfiguration, characterized by redistribution of edges from backbone to non-backbone units leading to loss of hubs and less-interconnected communities. Re-exposure after chronic ketamine treatment no longer silenced population bursting, indicating tolerance. Together, these results show that ketamine acutely silences human organoid networks by disconnecting backbone units, while chronic exposure induces tolerance to re-silencing while reducing the number of backbone units and leaving the network less active and less connected. The organoid-MEA platform provides a scalable, human-relevant system for dissecting circuit-level drug effects.

## Introduction

Ketamine is a cyclohexanone derivative originally developed as a safer alternative to the N-methyl-D-aspartate (NMDA) receptor antagonist phencyclidine (PCP). When used as an anesthetic, PCP produced frequent postoperative psychoses and dysphoria, limiting its clinical use. Ketamine, a structural analog with approximately one-tenth the potency of PCP, was found to be safe and effective for clinical anesthesia, producing rapid analgesia and a characteristic “dissociative” state with relatively short duration and fewer severe emergence reactions ([1], [2]). In 2000, low-dose ketamine was reported to exert rapid and robust antidepressant effects in patients with major depression, sparking intense interest in its use as a treatment for mood disorders (e.g. [3]). Since then, ketamine has been transformative in psychiatry: S-ketamine (esketamine, Spravato®) has been approved by the U.S. FDA for treatment-resistant depression, and racemic ketamine is widely used off-label for similar indications.

Mechanistically, ketamine is best known as a non-competitive NMDA receptor antagonist, with low micromolar affinity and stereoselectivity that parallels its anesthetic actions [4]. However, its actions are considerably more complex. At the low doses used in psychiatry, NMDA receptor blockade disproportionately affects GABAergic interneurons, leading to disinhibition of glutamatergic pyramidal cells, a transient glutamate surge, and enhanced AMPA receptor signaling [5]. Downstream, AMPA activation engages mTORC1 and increases the production of neurotrophic factors, most notably brain-derived neurotrophic factor (BDNF), which has been implicated in ketamine’s sustained antidepressant effects [6]. In addition, ketamine interacts with cholinergic, monoaminergic (dopamine and serotonin), and opioid systems, and inhibits HCN1 channels, all of which likely contribute to its diverse clinical profile and complex neurophysiological signatures [7]. Recent work has also highlighted roles for neuromodulatory pathways such as adenosine signaling in prefrontal and hippocampal circuits [8].

Stereochemistry adds an additional layer of complexity. The (S)-enantiomer, esketamine, displays roughly three-fold greater analgesic potency and higher anesthetic efficacy than (R)-ketamine (arketamine), and the two enantiomers may differ in both therapeutic and side-effect profiles ([9]). Ketamine’s system-level actions are also region-specific: for example, suppression of burst firing in the lateral habenula has been implicated in its rapid antidepressant effects ([10]). Thus, ketamine’s actions emerge from a rich interplay of receptor-level pharmacology, cell-type specificity, and circuit context, making it challenging to dissect how a single drug produces such rapid and profound changes in human brain network dynamics.

Given this complexity, there is a growing need for reduced yet human-relevant experimental systems to bridge molecular mechanisms and circuit-level population activity. Human brain organoids derived from induced pluripotent stem cells provide a scalable model that recapitulates key aspects of human cortical development and physiology. When coupled with high-density microelectrode arrays (MEAs), organoids exhibit spontaneous, structured burst dynamics and network-level oscillatory activity that approximate some features of human brain physiology (e.g. [11], [12]). This organoid–MEA platform has been recognized by regulators, including the U.S. FDA, as part of a broader transition toward human cell–based systems for pharmacological testing and safety evaluation ([13]).

In this study, we leverage human dorsal forebrain organoids interfaced with high-density MEAs to characterize how ketamine reshapes human neuronal network dynamics across multiple scales. We first quantify the acute effects of ketamine on population bursting and spike-sorted single-unit firing and inter-spike interval analysis. To link these changes to network-level organization, we compute pairwise functional connectivity and construct weighted functional graphs from which we show changes in standard graph-theoretic measures induced by ketamine. Finally, we compare acute and chronic exposure paradigms to determine how prolonged ketamine treatment alters subsequent sensitivity to the drug. Together, this approach combines human organoid models, high-density recordings, and network analysis to provide a systems-level view of ketamine’s actions on human cortical-like circuits.

## RESULTS

### Ketamine abolishes population bursting in cortical organoids by disrupting network connectivity

To investigate the effects of acute ketamine dose on human brain organoids, we plated 14 six-month-old dorsal forebrain organoids [14] on multi-electrode arrays (MEAs)[15]. First, we recorded baseline activity for 5 minutes, during which time all 14 organoids displayed population bursting with an average inter-burst interval of 10 ± 4.4 s. Bursts were defined as population activity smoothed with an average sliding window of 200 ms exceeding 3 sigma (Methods).

Next, we added ketamine to the media to reach a final concentration of 20 μg/mL, added an equal volume of saline to the control group, and resumed recording immediately. Ketamine treatment rapidly and consistently abolished population bursting (n = 8; Figure 1A, Supp. Figure 1A), whereas saline had no effect (n = 6; Figure 1B, Supp. Figure 1B). Spike sorting with Kilosort4 yielded an average of 90 ± 30 units per organoid. The number of active units was unchanged in four organoids; two organoids lost one unit each, and one organoid lost two units, indicating that the abolition of bursting is not explained by unit silencing alone. The average firing rate per organoid decreased following ketamine treatment (Figure 1C); however, only a subset of individual units exhibited reduced firing rates (Figure 1D). Due to inherent fluctuations in firing rates, some units showed reduced activity following both saline and ketamine treatment; however, there are significantly more units exhibiting a ≥25% decrease in firing rate after ketamine treatment compared to saline (Figure 1E). Comparable proportions of units showed a ≥25% increase in firing rate after saline and ketamine treatment (∼10%), indicating that ketamine does not specifically elevate firing activity. The firing rate reductions were not confined to the bursts; firing rates in the inter-burst intervals also decreased (Supp. Figure 2A).

**Figure 1.**
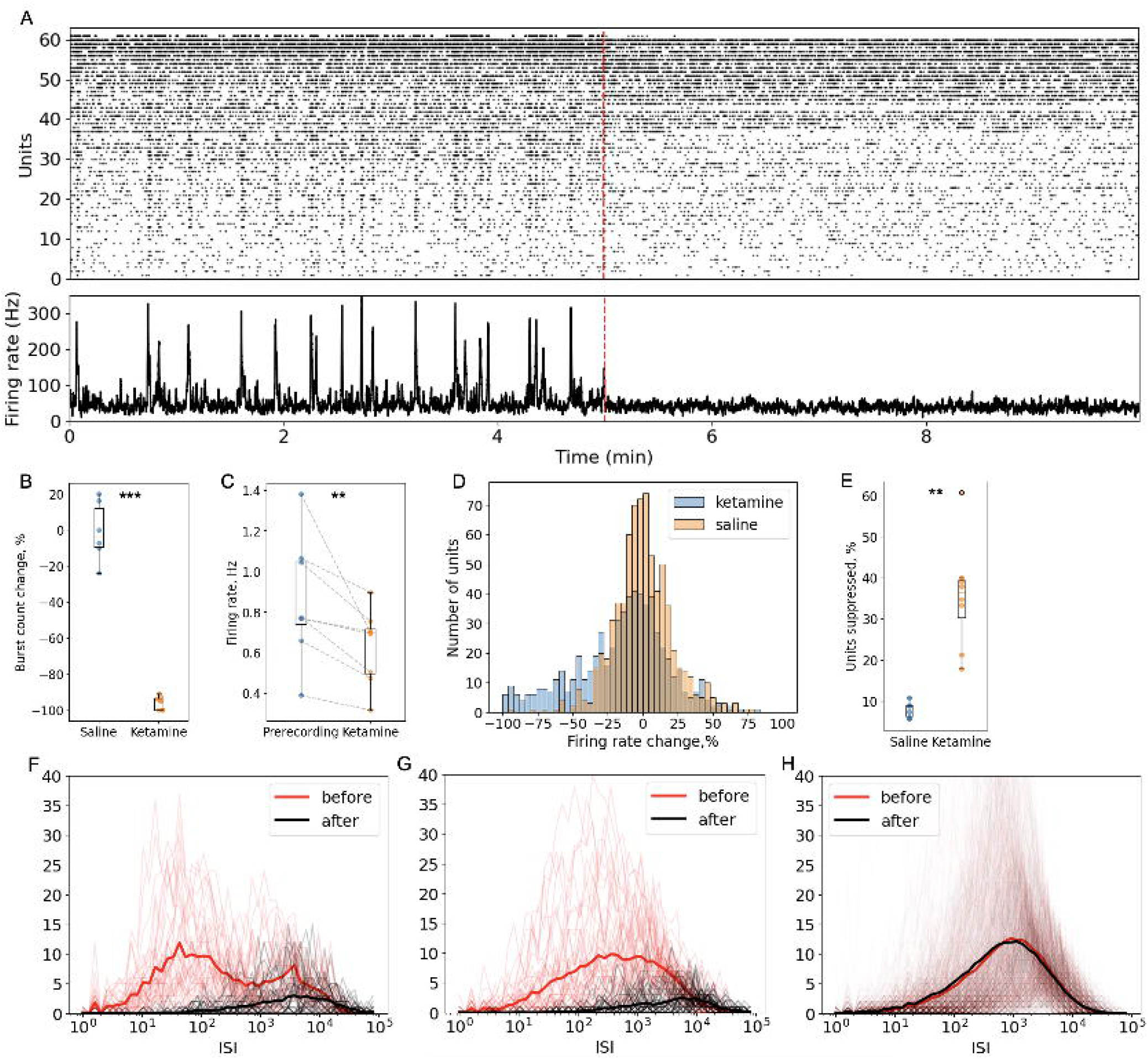
Acute ketamine abolishes bursting and reshapes single-unit activity in human forebrain organoids. A. Representative MEA/raster traces showing abolition of bursting after ketamine. B. Percent change in burst count with saline vs ketamine treatment. C. Mean firing rate per organoid before and after treatment. D. Distribution of unit-wise firing-rate changes with ketamine vs saline treatment. E. Proportion of units with ≥25% firing-rate decrease with ketamine vs saline. F. ISI distributions for bimodal units, demonstrating loss of the <100 ms “fast” mode after ketamine. G-H. ISI distributions for unimodal units, illustrating shifts toward longer intervals in affected units (G).

We identified a subgroup of units that consistently participate in bursts. Such units were previously identified in organoids at baseline conditions as “backbone” units ([11]). Here we define a backbone unit as firing at least twice in ≥85% of bursts, yielding a total of 15% of all recorded units. Backbone units are characterized by increase in firing rate during bursts, and a slightly elevated baseline activity during inter-burst intervals as compared to non-backbone units (Figure 2A, B, Supp. Figure 2C). Moreover, backbone units started firing earlier than non-backbone units within bursts (Figure 2C), which implicates them as drivers of burst initiation.

**Figure 2.**
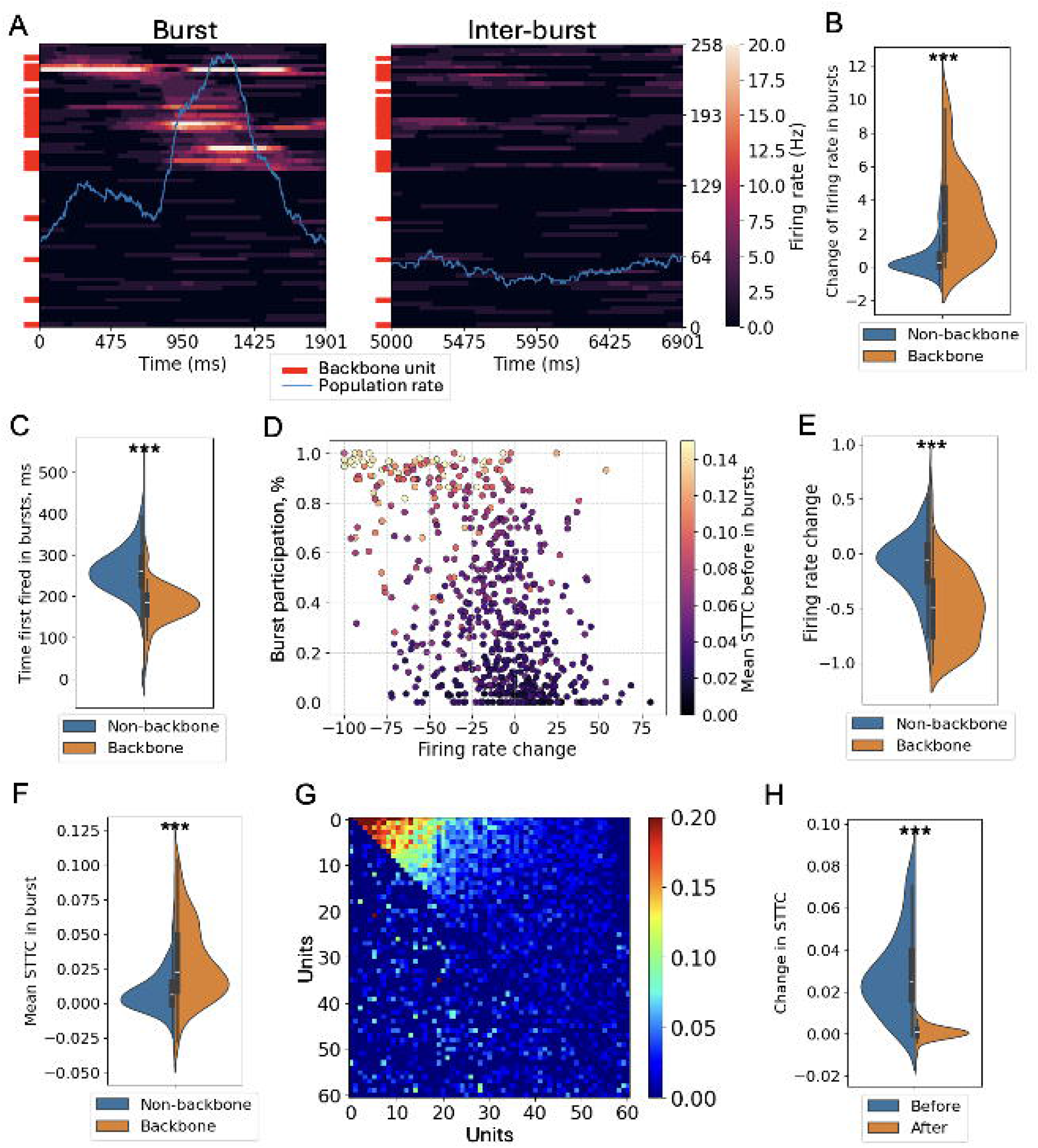
Ketamine reduces functional connectivity and selectively silences “backbone” units in human forebrain organoids. A. Example of a zoomed in burst and an inter-burst interval. Brighter colors reflect higher firing rates of individual units, blue line shows population firing rate. Red ticks indicate backbone units. B. Burst-evoked change in firing rate for backbone versus non-backbone units. C. Backbone units fire earlier in bursts than non-backbone. D-E. Preferential firing rate suppression of backbone units by ketamine expressed in % firing change (ΔFR, refer to Methods). F. Functional connectivity (STTC) during bursts for backbone versus non-backbone units. G. Pairwise STTC matrix before (upper triangle) and after (lower triangle) ketamine, showing a global reduction in STTC. H. Dramatic reduction in mean STTC per backbone unit with ketamine treatment.

Next, we examined inter-spike interval (ISI) distributions for each unit in each of 8 organoids treated with ketamine (Supp. Figure 3). Because bursts are characterized by rapid firing rates, most of the fast ISIs (<100 ms), approximately 56%, occurred within bursts. Among a total of 613 units, 53 displayed bimodal ISI distributions, of which 51 lost the “fast” mode following ketamine treatment, essentially abolishing fast ISIs (Figure 1F). The remaining units exhibited unimodal ISI distributions that shifted toward longer intervals in affected units, while the majority were unaffected by ketamine (Figure 1G, H).

Importantly, backbone units were preferentially suppressed by ketamine, in contrast to saline-treated controls (Figure 2D, E, Supp. Figure 2B), suggesting a potential mechanism for ketamine-induced burst loss. Ketamine selectively reduced the firing rate of backbone units not only during bursts but also in inter-burst periods (Supp. Figure 2D); however, their activity was not abolished entirely (Supp. Figure 2E).

To quantify how ketamine treatment affects functional connectivity between units, we measured pairwise coupling with the spike-time tiling coefficient (STTC) which measures the degree of temporal correlation between spike trains while correcting for differences in firing rate [16]. Backbone units exhibited high connectivity, as reflected by an elevated STTC (Figure 2F) both in bursts and inter-burst regions, compared to non-backbone units Supp. Figure 2F).

Ketamine treatment produced a marked reduction in STTC, evident in pairwise matrices sorted by pre-treatment connectivity (Figure□2G) and absent with saline (Supp. Figure 2G). Backbone units became functionally disconnected from the network with ketamine treatment, leading to a significant reduction in STTC among these units overall (Figure 2H) and during inter-burst intervals (Supp. Figure 2H). Together, these results indicate that ketamine’s population-level silencing effect is mediated by selective suppression and disconnection of backbone units, which normally drive burst activity.

We constructed functional connectivity graphs for each organoid before and after ketamine treatment (Figure 3A), representing units as nodes and significant functional connections as weighted edges (defined as STTC values exceeding chance levels, *p* < 0.05; see Methods). The number of such edges decreased significantly with ketamine treatment (Figure 3B) along with the average edge weight (Figure 3C). Consistent with backbone units serving as drivers of population activity, they exhibited a significantly higher proportion of hub nodes (defined as units with >5 edges) compared to non-backbone units (Figure 3D), forming a dense, interconnected network around them. Ketamine treatment decentralized the network, markedly reducing the total number of hub nodes, eliminating them entirely in several organoids (Figure 3E). The concurrent increase in modularity (the extent to which nodes cluster into densely connected communities) (Figure 3F) confirms that the network becomes more segregated, with nodes clustering into more distinct, less-interconnected communities. This is further supported by reductions in average clustering, global efficiency (the average inverse shortest-path length, reflecting how easily any node can be reached from any other), and transitivity (the fraction of closed triangles in the network, reflecting triangle-based local clustering) (Figures 3G–I), all of which point to a breakdown in local and global communication efficiency. Together, these findings suggest that ketamine shifts the network toward a more fragmented and less integrated topology, potentially impairing coordinated information flow across the network while promoting isolated local activity. Such reconfiguration may reflect ketamine’s known impact on disrupting large-scale cortical integration, possibly underlying some of its dissociative and cognitive effects.

**Figure 3.**
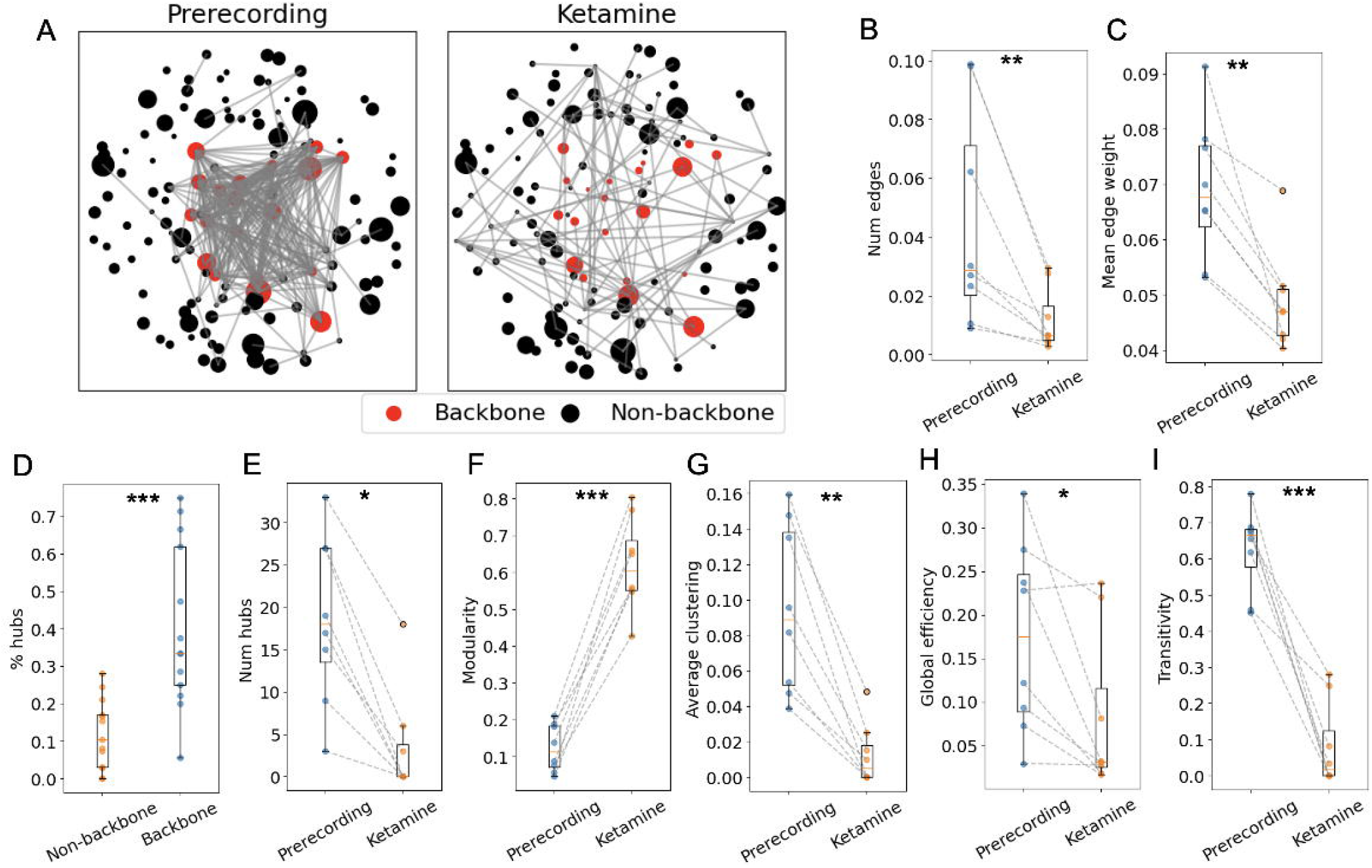
Ketamine decentralizes organoid networks and shifts it toward a more fragmented state. A. Functional connectivity graphs pre- and post-ketamine. B. Number of significant edges per organoid before vs after ketamine. C. Mean edge weight (STTC) before vs after ketamine. D. Proportion of hub nodes (>5 edges) among backbone vs non-backbone units. E. Total hub count per organoid before vs after ketamine. F. Modularity before vs after ketamine. G - I. Average clustering coefficient, transitivity, and global efficiency before vs after ketamine.

### Chronic exposure to ketamine attenuates population burst silencing upon re-exposure

To investigate the effects of chronic ketamine exposure on organoid network activity, organoids were continuously treated with ketamine for 6 days and then transferred to a regular culture medium on day 6 (“chronic protocol”). Control conditions included (i) organoids given matched saline volumes after the initial ketamine exposure (“acute protocol”) and (ii) organoids exposed only to saline (“control protocol”). On day 7, a 10-min baseline recording was obtained, after which organoids were re-exposed to 20 μg/mL ketamine. The protocols are summarized in the Figure 4A. Baseline recordings indicated that bursting had resumed, and re-exposure no longer suppressed network activity after the chronic protocol (Figure 4B). In contrast, re-exposure after the acute protocol silenced bursts again, recapitulating the primary exposure effect (Figure 4C).

**Figure 4.**
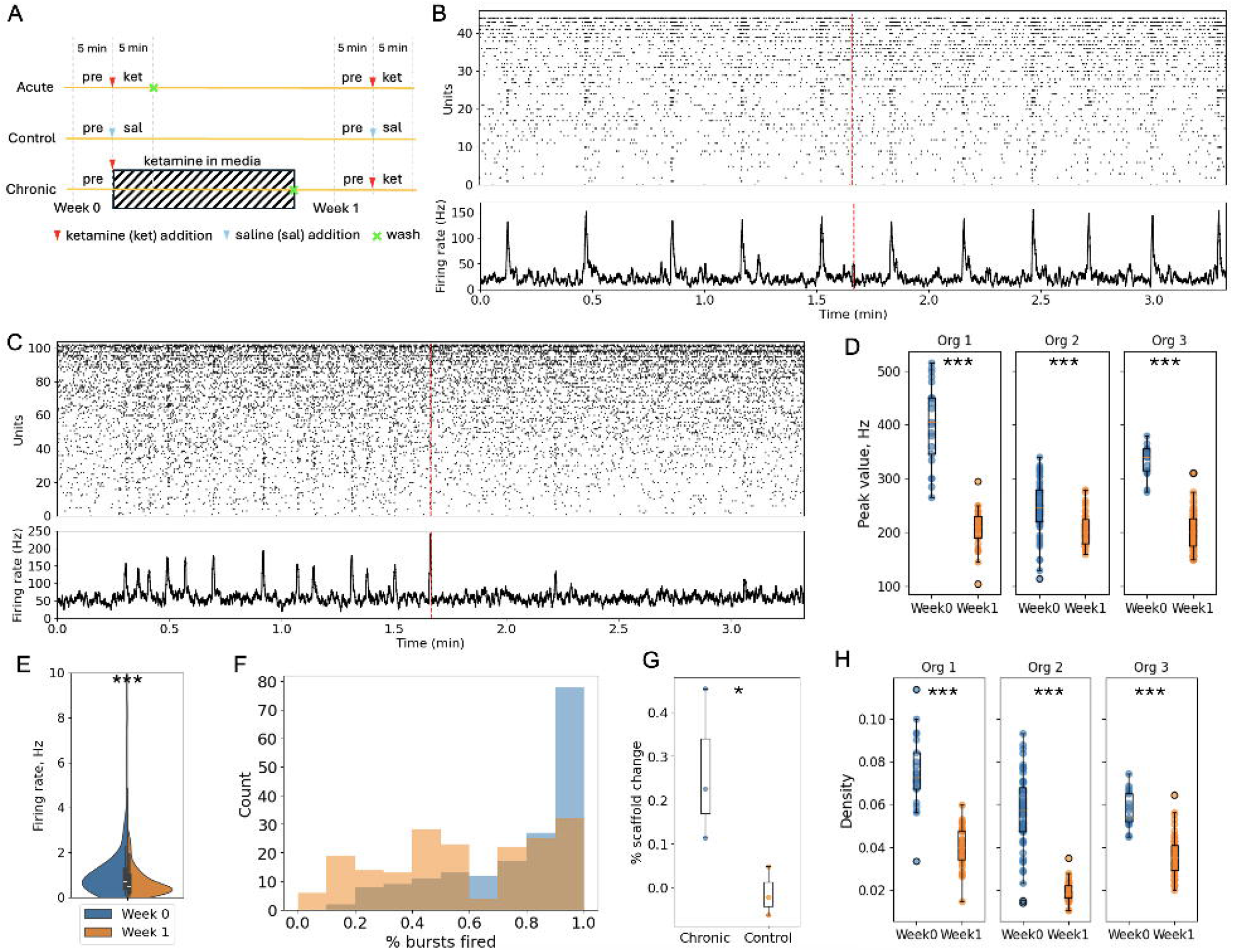
Chronic ketamine induces tolerance to re-exposure and reduces baseline correlation. A. Schematic illustration of the chronic and acute ketamine exposure protocols and the corresponding control condition. B. Representative MEA/raster traces of an organoid chronically exposed to ketamine for a week: bursting activity has recovered and is no longer suppressed by ketamine addition. C. Re-exposure to ketamine a week after a one-time acute exposure has the same burst silencing effect as on day 0. D. Baseline burst intensity is significantly reduced in each of the three organoids chronically exposed to ketamine for a week. E. Baseline firing rate distribution is shifted towards lower values after chronic ketamine exposure. F-G. Proportion of units consistently participating in bursts decreases with chronic ketamine exposure. H. Within-burst connectivity at baseline is reduced after chronic ketamine exposure, leading to significantly less percent of significant edges (reduced graph density).

Ketamine tolerance following chronic exposure was further evidenced by the absence of preferential firing-rate suppression of backbone units and by unchanged connectivity upon re-exposure (Supp. Figure 2I-K). Nevertheless, the baseline after chronic exposure differed markedly from the initial pre-exposure baseline. Maximum population activity during burst (Figure 4D) and individual unit firing rates (Figure 4E) were significantly reduced. The proportion of units that consistently participated in bursts - including backbone units - also declined (Figure 4F, G). Consistent with these changes, within-burst connectivity decreased as well, showing a significant reduction in graph density (Figure 4H). Together, these findings indicate that chronic ketamine exposure weakens the network while still permitting the emergence of population bursts.

### Gradual ketamine dose increase reveals nonlinear suppression and partial recovery of population bursts in organoids

Finally, we quantified network dynamics during a gradual increase of ketamine dose. We recorded neuronal activity from the same organoid (n=2) throughout sequential ketamine additions (eight consecutive steps from 0 to 40 µM), followed by a 24 h continuous exposure at the final high dose (40 µM), and then a recovery phase in which ketamine was washed out and replaced with saline for 24 h before re-recording (Figure 5A, for individual raster plots refer to Supp. Figure 4). Across the dose series, network bursting was highly sensitive to ketamine: total burst counts dropped dramatically after the first low-dose addition (2.5 µM), followed by a partial rebound and re-emergence of bursting at subsequent doses and complete burst silencing starting from 15 µM dose (25 µM in the second organoid, Figure 5B, Supp. Figure 5A, B).

**Figure 5.**
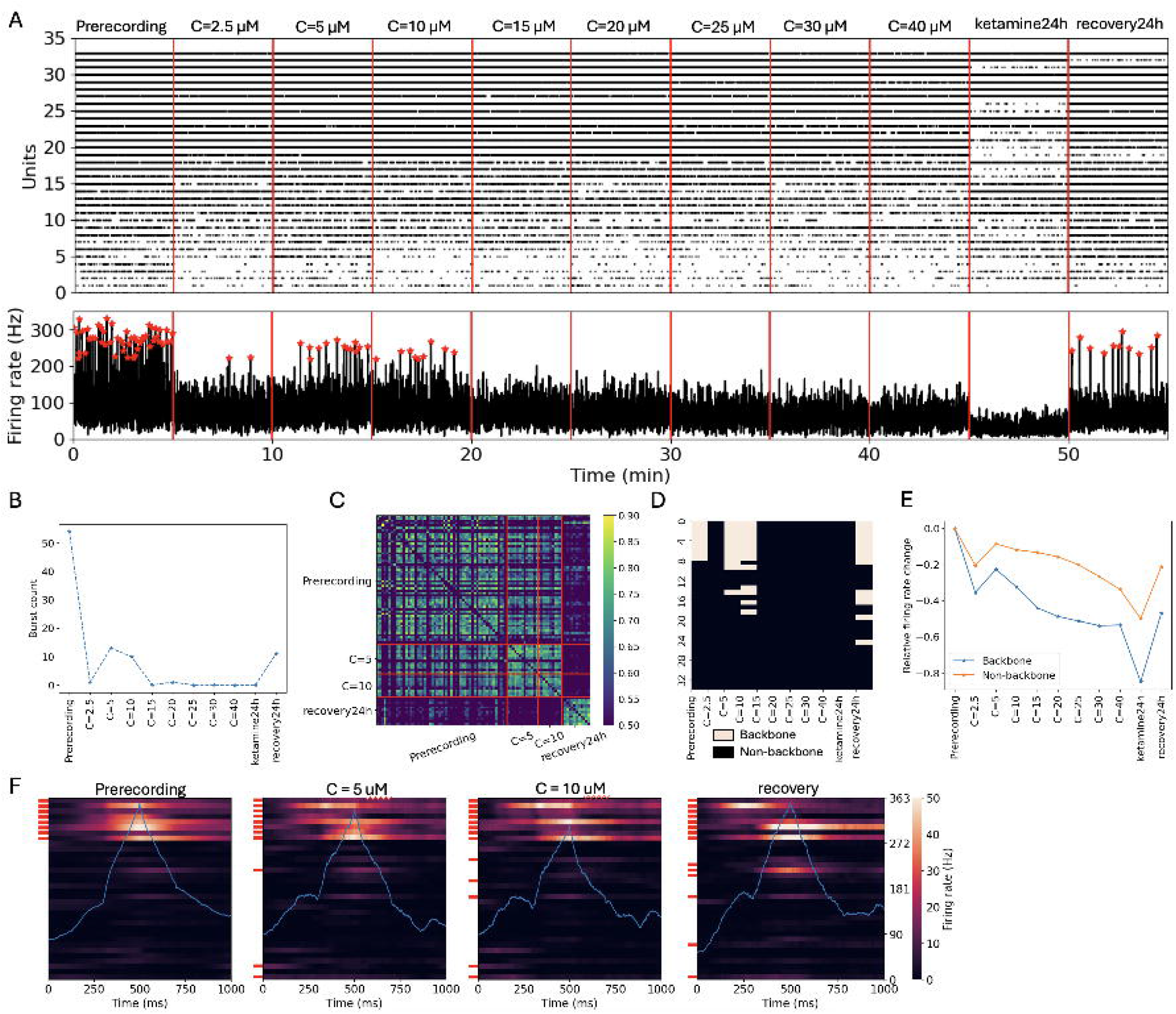
Ketamine dose–response and recovery. (A) Raster plot from a single organoid during eight consecutive ketamine dose steps (0–40 μM), followed by 24 h of continuous high-dose exposure and a subsequent 24 h recovery period in saline. (B) Total burst count as a function of dose. Bursting drops sharply after the first ketamine addition (2.5 μM), partially rebounds at intermediate doses, and returns after 24 h recovery in saline. (C) Burst-to-burst firing-rate cross-correlation indicates that burst structure does not return to its pre-exposure pattern after 24 h recovery. (D) Backbone units identified during the baseline recording largely remain backbone units throughout the dose–response protocol; additional backbone units emerge with increasing dose and after recovery. (E) Relative changes in mean firing rate show that backbone units reduce firing more rapidly than non-backbone units as dose increases and fail to fully recover after the saline recovery period. (F) Average raster plot of bursts at different ketamine concentrations and during recovery. Red bars indicate scaffolds. Blue line indicates average population rate during burst.

Prolonged high-dose exposure produced profound spiking suppression. After washout and 24 h in saline, bursting activity returned, indicating substantial recovery of gross network activity. However, this recovery incompletely restored the temporal organization: burst-to-burst firing-rate cross-correlations showed that bursts did not regain their pre-exposure structure after 24 h in saline (Figure 5C, Supp. Figure 5C). On average, bursts gradually changed their temporal structure with dose increase and did not regain their baseline pattern after recovery (Figure 5F). Tracking backbone units revealed further persistence and remodeling: units identified as backbones at baseline largely remained backbones throughout the dose-response, while additional backbone units emerged with increasing dose and after recovery (Figure 5D, Supp. Figure 5D). Consistent with this, backbone units exhibited a faster dose-dependent reduction in mean firing rate than non-backbone units and firing of both groups did not fully recover after the 24 h saline recovery period (Figure 5E, Supp. Figure 5E).

## Discussion

This study uses human dorsal forebrain organoids coupled to MEAs to bridge single-unit physiology, functional connectivity, and network organization in response to ketamine. Three main findings emerge. First, an acute ketamine dose rapidly abolishes population bursting while the number of active units remains largely unchanged; firing rates decline only in a subset of units. This result shows that acute ketamine extinguishes bursts primarily by disrupting temporal synchrony, not by wholesale unit silencing. We show that ketamine preferentially suppresses and functionally disconnects “backbone” units, units that are enriched for hub-like properties at baseline - higher connectivity, earlier onset within bursts, and greater burst-evoked rate increases - suggesting they significantly contribute to initiating and sustaining coordinated events. Together, these observations provide a parsimonious mechanism: ketamine’s selective suppression and disconnection of backbone units causes the loss of bursting.

Second, this loss of bursting is accompanied by a marked reduction in functional coupling and a reorganization of network topology - fewer and weaker connections, loss of hubs, increased modular segregation, and diminished local and global efficiency. In other words, ketamine drives a shift toward sparser, more modular graphs with reduced clustering and efficiency - features consistent with fragmentation and impaired information flow. In cortical-scale terms, such a reconfiguration resembles a move from an integrated regime toward segregated processing, aligning conceptually with ketamine’s dissociative state.

Finally, the chronic-exposure results refine this picture. After six days of ketamine followed by washout, organoids recovered bursting and no longer showed re-silencing upon re-exposure suggesting development of tolerance. However, the recovered baseline was not a simple reversion: burst intensity, mean firing rate, backbone participation, and within-burst connectivity all remained reduced. One interpretation is that homeostatic and plastic mechanisms - synaptic scaling, receptor trafficking, or network rewiring - reconfigure the circuit to restore the capacity for bursts without reinstating the previous reliance on strongly connected backbone units. In practical terms, the network becomes less susceptible to ketamine’s acute silencing while operating in a lower-gain, less-integrated state. Such decoupling of “ability to burst” from “high integration and hubness” may help explain why repeated ketamine exposure *in vivo* can show tolerance to certain acute electrophysiological signatures while leaving enduring alterations in network dynamics.

Notably, in a within-organoid dose–response experiment, bursting collapsed after the first low-dose ketamine addition (2.5 µM) but then partially rebounded during subsequent dose steps, consistent with rapid compensatory plasticity and network remodeling even on the timescale of a single session. After a prolonged 24 h exposure at 40 µM, activity was near-completely silenced; nevertheless, bursts re-emerged only after ketamine washout and a full 24 h in saline, indicating that gross bursting capacity can recover within 24 h while finer-grained burst structure and backbone-unit firing show incomplete restoration.

The cortical organoid-MEA platform proved sensitive to both acute and chronic drug effects across scales - from ISIs to graph topology - and revealed subpopulation-specific vulnerabilities not apparent from averages alone. This multi-dimensional readout enables discriminative extraction of network features (bursting, ISIs, coupling, hubness, modularity, efficiency) for comparative pharmacology, screening, and safety assessment. Limitations include that STTC-based functional connectivity explores temporal correlations rather than synaptic connections and can reflect shared inputs or rate changes despite controls. Additionally, organoids lack vascularization, full cell-type diversity, and subcortical structures, including the thalamus - a key regulator of sleep-wake state and thalamocortical rhythms - so state-dependent dynamics and ketamine effects that rely on corticothalamic interactions cannot be modeled here. Future work will expand dose-time mapping, perform enantiomer-specific tests, and integrate single-cell transcriptomics. In summary, acute ketamine extinguishes bursting by selectively suppressing and disconnecting hub-enriched “backbone” units, fragmenting connectivity, whereas chronic exposure confers tolerance to re-silencing yet leaves a persistently lower-activity, less-integrated baseline. Thus, organoid–MEA assays provide a human-relevant platform to resolve multi-scale circuit pharmacology and to link therapeutic actions with network-level liabilities.

## METHODS

### Human brain organoid generation

The human induced pluripotent stem cell (iPSC) line F12442.4 was provided by Karch [17]. Cells were maintained in mTeSR Plus medium (STEMCELL Technologies) on Matrigel-coated (Corning) 6-well plates with daily medium changes and passaged using ReLeSR (STEMCELL Technologies) at 70–80% confluency. Human dorsal forebrain organoids were generated using established protocols [18–20]. On day 0, hiPSCs were dissociated into single cells with Accutase (STEMCELL Technologies) and seeded into 96-well “slit-well” plates (S-Bio) at 1 × 10□ cells per well in mTeSR Plus supplemented with 10 nM ROCK inhibitor. After 24 h, cultures were switched to Neural Induction Medium (NIM) composed of Essential 6 (Gibco) with 1× Antibiotic–Antimycotic (Gibco), 2.5 µM dorsomorphin, 10 µM SB-431542, and 2.5 µM XAV-939 (all from Tocris). Dorsal forebrain organoids were fed daily with NIM from days 1–5.

On day 6, organoids were transferred to Neural Differentiation Medium (NM) containing Neurobasal-A (Gibco), B-27 without vitamin A (Gibco), 1× GlutaMAX (Gibco), and 1× Antibiotic–Antimycotic, supplemented with 20 ng/mL EGF and 20 ng/mL FGF2 (Shenandoah). NM with EGF and FGF2 was changed daily from days 6–15 and every other day from days 16– 24.From days 25–42, NM was supplemented with 20 ng/mL BDNF and 20 ng/mL NT3 (Shenandoah), with medium changes every other day. From day 43 onward, organoids were maintained in NM without additional growth factors and fed every four days. Organoids were used for electrophysiological experiments at 4–7 months of differentiation.

### Organoid interfacing with CMOS microelectrode arrays (MEAs)

Prior to organoid placement, the recording surfaces of CMOS MEAs (MaxOne, Maxwell Biosystems) were coated with poly-L-lysine (PLL) and laminin. Briefly, 0.5 mL of PLL solution (0.1 mg/mL in ultrapure water; Sigma-Aldrich) was added to each MEA reservoir and incubated for ∼1 h at 37 °C. Arrays were then transferred to a sterile hood, the PLL solution was aspirated, and the wells were washed three times with 1 mL ultrapure water. The wells were allowed to dry for ∼1 h under sterile conditions. Next, 0.5 mL laminin solution was added to each reservoir and incubated for ∼3 h at 37 °C. The laminin solution was then aspirated and the wells were washed three times with ultrapure water as above.

Whole organoids were transferred to the CMOS MEA well using a cut P1000 pipette tip and gently positioned over the recording electrode area with sterile forceps. Organoids were allowed to adhere to the surface for ∼1 min, after which a 50 µL drop of BrainPhys medium (STEMCELL Technologies) was gently added on top of each organoid. The MEAs were returned to the incubator (5% CO□, 37 °C) for ∼1 h, after which 0.5 mL of BrainPhys was added to each reservoir. Organoids on MEAs were maintained in the incubator (5% CO□, 37 °C), and the medium was exchanged twice per week for two weeks prior to recording.

### Electrophysiology recordings and pharmacology

Whole brain organoids were recorded at 4–7 months of differentiation (see Section 1). High-density extracellular field potentials were measured inside a cell culture incubator (5% CO□, 37 °C) using CMOS microelectrode array technology (MaxOne, Maxwell Biosystems, Zurich, Switzerland). Each sensor contained 26,400 recording electrodes (7.5 µm diameter, 17.5 µm center-to-center spacing) within a sensing area of 3.85 × 2.10 mm. Of these, up to 1,024 electrodes could be selected for simultaneous recording. On-board low-noise amplifiers included a 0.5 Hz high-pass filter to minimize slow drift. Automatic activity scans (tiled blocks of 1,020 electrodes) were performed to identify the spatial distribution of electrical activity across the organoid surface. All recordings were sampled at 20 kHz and saved in HDF5 format.

For detailed recordings, we selected the 1,020 electrodes with the highest spiking activity while maintaining a minimum spacing of at least two electrode pitches (2 × 17.5 µm). This configuration provided sufficient electrode redundancy per neuron to enable accurate identification of single units by spike sorting while still resolving network activity across the entire organoid.

Racemic ketamine (Sigma-Aldrich) was reconstituted to a stock concentration of 100 mg/mL in PBS and further diluted to 1 mg/mL aliquots in PBS. During each experiment, baseline activity was first recorded, after which 20 µL of the ketamine aliquot was added to 0.5 mL of medium in the recording well, yielding a final concentration of 20 µg/mL (scaled proportionally for other concentrations). For control conditions, 20 µL of PBS was added instead. Recordings were immediately resumed (or continued) following ketamine or saline addition and continued for an additional 5 min.

### Spike sorting

Raw extracellular recordings were automatically spike sorted to extract single-unit activity using the Kilosort 4 algorithm [21]. For recordings obtained on the same day from the same chip, continuous traces were concatenated in time prior to spike sorting so that units could be tracked across blocks within a day. Putative units underwent manual curation by visual inspection of voltage traces on electrodes surrounding each unit’s spatial peak, as well as examination of waveform shape and stability over time. Units with mean firing rates below 0.083 Hz (fewer than 5 spikes per minute over the recording) were excluded from further analysis.

### Firing rate and inter-spike interval analysis

For each unit and recording, the firing rate (Hz) was computed as the total number of spikes divided by the recording duration in seconds. Changes in firing rate between two recordings (e.g. before and after treatment) were quantified using a symmetrized difference:

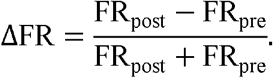

Inter-spike intervals (ISIs) were defined as the time differences between consecutive spikes from the same unit. To classify the modality of ISI distributions, we used a combined model-based and density-based approach in log-ISI space. For each unit, we fit Gaussian mixture models with one and two components and compared their Bayesian Information Criterion (ΔBIC = BIC□ − BIC□). In parallel, we estimated a kernel density of the ISI distribution and counted the number of peaks. ISI distributions were labeled as bimodal if there was strong evidence for two components (ΔBIC ≥ 6) and at least two peaks in the density estimate; otherwise they were labeled unimodal. For bimodal units, the split point between modes was defined as the local minimum of the density between the two mixture means and reported in the original time units.

### Burst detection

Bursts were defined from the population spiking activity. First, we constructed a population activity trace by summing spikes across all recorded units in a sliding window of 200 time bins (bin width 1 ms). The resulting signal was smoothed with a Gaussian kernel (σ = 50 bins) and baseline-corrected by subtracting its mean. Putative bursts were identified as local maxima in this trace whose amplitudes exceeded three standard deviations above the baseline-corrected mean. For each detected peak, burst onset and offset were defined as the time points at which the baseline-corrected trace crossed 5% of that peak’s amplitude on the rising and falling edges, respectively.

Backbone units were defined as units that fired at least two spikes in ≥85% of detected bursts, resulting in approximately 15% of all recorded units being classified as backbone. All other units were considered non-backbone.

### Spike time tiling coefficient (STTC) and functional connectivity graphs

Pairwise functional coupling between units was quantified using the spike time tiling coefficient (STTC), which measures the degree of temporal correlation between two spike trains while correcting for differences in firing rate [22]. For each recording and condition, we computed STTC for all pairs of simultaneously recorded units using a coincidence window of Δt = 20 ms. STTC was computed both for full recordings and, where indicated, separately within burst and inter-burst periods using the burst definitions above.

To assess statistical significance, we generated null distributions of STTC values from shuffled spike rasters. For each recording, surrogate datasets were created by repeatedly selecting two units at random, choosing one spike time from each, and swapping these spike times between the units. This spike-swap procedure was iterated 500 times per recording, preserving spike counts per unit and overall burst structure while disrupting precise pairwise temporal structure. For each unit pair, the observed STTC was compared to the distribution of STTC values obtained from the shuffled rasters, and a one-sided p-value was defined as the proportion of surrogate STTC values greater than or equal to the observed value. Pairwise connections were considered significant if p < 0.05.

For each organoid and condition, we constructed a functional connectivity graph in which nodes represented single units and undirected, weighted edges represented significant functional connections. The weight of each edge was given by the observed STTC value for that unit pair, and edges that did not reach the p < 0.05 threshold were omitted (weight set to zero). This yielded a weighted adjacency matrix for each organoid before and after treatment.

Graph-theoretic properties were computed on these adjacency matrices using the NetworkX Python library. Node degree was defined as the number of suprathreshold edges incident on each node; units with degree > 5 were classified as hub nodes. Network density was computed as the fraction of present edges out of all possible undirected edges. Where reported, community structure was quantified by modularity using a Louvain-type algorithm, local structure by the weighted clustering coefficient and transitivity, and global communication by global efficiency (the average inverse shortest path length between all node pairs in the weighted graph).

## Acknowledgements

We thank Tiny Blue Dot Foundation for contributing to the support of this work. We thank Max Lim and Tjitse Van der Molen for technical support on running the spike sorting.

## Notes

### Competing Interest Statement

The authors have declared no competing interest.

## References

1. Domino EF, Chodoff P, Corssen G. Pharmacologic effects of CI-581, a new dissociative anesthetic, in man. Clin Pharmacol Ther. 1965;6(3):279–291.

2. Corssen G, Domino EF. Dissociative anesthesia: further pharmacologic studies and first clinical experience with the phencyclidine derivative CI-581. Anesth Analg. 1966;45(1):29–40.

3. Berman RM, Cappiello A, Anand A, Oren DA, Heninger GR, Charney DS, et al. Antidepressant effects of ketamine in depressed patients. Biol Psychiatry. 2000;47(4):351–354.

4. Zorumski, C. F., Izumi, Y., & Mennerick, S. Ketamine: NMDA receptors and beyond. J. Neurosci. 2016; 36(44), 11158–11164. doi:10.1523/JNEUROSCI.1547-16.2016

5. Homayoun H., Moghaddam B. NMDA receptor hypofunction produces opposite effects on interneurons and pyramidal neurons. J. Neurosci, 2007; 27, 11496–11500. doi:10.1523/JNEUROSCI.2213-07.2007

6. Li N., Lee B., Liu R., Banasr M., Dwyer J. M., Iwata M., Li X., Aghajanian G., Duman R. S. mTOR-dependent synapse formation underlies the rapid antidepressant effects of NMDA antagonists. Science. 2010; 329, 959–964. doi:10.1126/science.1190287

7. Sleigh J., Harvey M., Voss L., Denny B. Ketamine – more mechanisms of action than just NMDA blockade. TACC. 2014; 4, 76–81. doi:10.1016/j.tacc.2014.03.002

8. Yue C., Wang N., Zhai H., et al. Adenosine signalling drives antidepressant actions of ketamine and ECT. Nature. 2025; 24, 20–30. doi:10.1038/s41586-025-09755-9

9. Sartorius A, Kranaster L. Ketamine, esketamine, and arketamine: their mechanisms of action and clinical applications in psychiatry. Biomedicines. 2024;12(10):2283. doi:10.3390/biomedicines12102283.

10. Yang Y, Cui Y, Sang K, Dong Y, Ni Z, Ma S, et al. Ketamine blocks bursting in the lateral habenula to rapidly relieve depression. Nature. 2018;554(7692):317–322. doi:10.1038/nature25509.

11. van der Molen T, Spaeth A, Chini M, et al. Preconfigured neuronal firing sequences in human brain organoids. Nat Neurosci. 2025; doi:10.1038/s41593-025-02111-0.

12. Sharf T, van der Molen T, Glasauer SMK, Guzman E, Buccino AP, Luna G, et al. Functional neuronal circuitry and oscillatory dynamics in human brain organoids. Nat Commun. 2022;13(1):4403. doi:10.1038/s41467-022-32115-4.

13. Zushin PJH, Mukherjee S, Wu JC. FDA Modernization Act 2.0: transitioning beyond animal models with human cells, organoids, and AI/ML-based approaches. J Clin Invest. 2023;133(21):e175824. doi:10.1172/JCI175824.

14. Pas□ca AM, Sloan SA, Clarke LE, Tian Y, Makinson CD, Huber N, et al. Functional cortical neurons and astrocytes from human pluripotent stem cells in 3D culture. Nat Methods. 2015;12(7):671–678. doi:10.1038/nmeth.3415.

15. Obien MEJ, Gong W, Frey U, Bakkum DJ. CMOS-based high-density microelectrode arrays: technology and applications. In: Emerging Trends in Neuro Engineering and Neural Computation. Singapore: Springer; 2017. p. 3–39. doi:10.1007/978-981-10-3957-7_1.

16. Cutts CS, Eglen SJ. Detecting pairwise correlations in spike trains: an objective comparison of methods and application to the study of retinal waves. J Neurosci. 2014 Oct 22;34(43):14288–303. doi: 10.1523/JNEUROSCI.2767-14.2014.

17. Karch CM, Hernández D, Wang JC, et al. Human fibroblast and stem cell resource from the Dominantly Inherited Alzheimer Network. Alzheimers Res Ther. 2018;10(1):69. doi:10.1186/s13195-018-0400-0.

18. Sloan SA, Andersen J, Pas□ca AM, Birey F, Pas□ca SP. Generation and assembly of human brain region–specific three-dimensional cultures. Nat Protoc. 2018;13(9):2062– 2085. doi:10.1038/s41596-018-0032-7.

19. Yoon SJ, Elahi LS, Pas□ca AM, Marton RM, Gordon A, Revah O, et al. Reliability of human cortical organoid generation. Nat Methods. 2019;16(1):75–78. doi:10.1038/s41592-018-0255-0.

20. Bertucci T, Bowles KR, Lotz S, Qi L, Stevens K, Goderie SK, et al. Improved protocol for reproducible human cortical organoids reveals early alterations in metabolism with MAPT mutations. bioRxiv. 2023; 2023.07.11.548571. doi:10.1101/2023.07.11.548571.

21. Pachitariu, M., Sridhar, S., Pennington, J. et al. Spike sorting with Kilosort4. Nat Methods. 2024; 21, 914–92. doi:10.1038/s41592-024-02232-7

22. Catherine S. Cutts and Stephen J. Eglen. Detecting Pairwise Correlations in Spike Trains: An Objective Comparison of Methods and Application to the Study of Retinal Waves. J. Neurosci. 2014; 34 (43) 14288–14303. doi:10.1523/JNEUROSCI.2767-14.

