## Supplemental Figures for "Neuronal Population Effects of Ketamine on Human Brain Organoids"

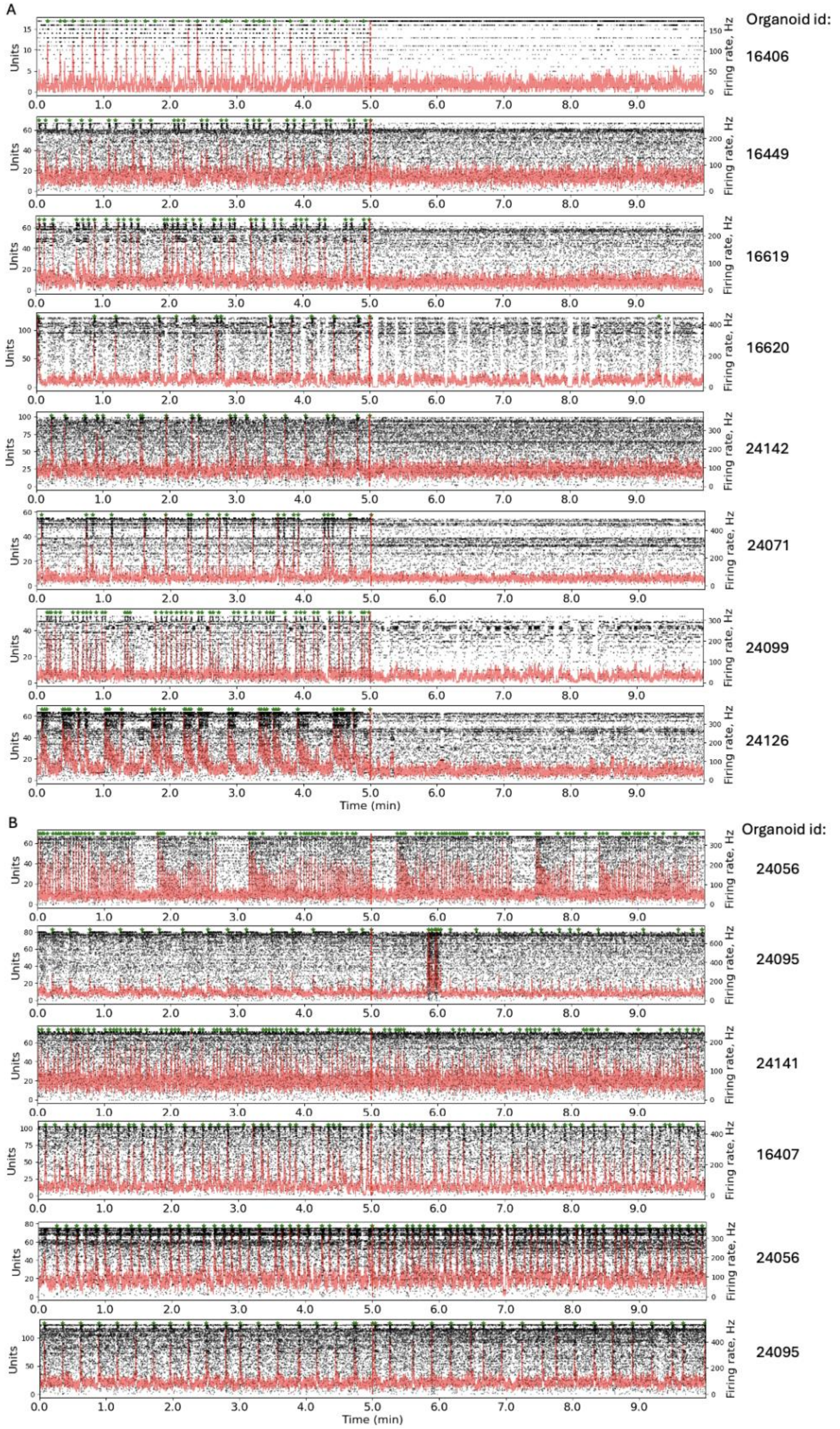


**Supplemental Figure 1.** A. Raster traces showing abolition of bursting after ketamine. Red dotted line indicates ketamine addition. B. Raster traces showing no abolition of bursting after saline treatment.

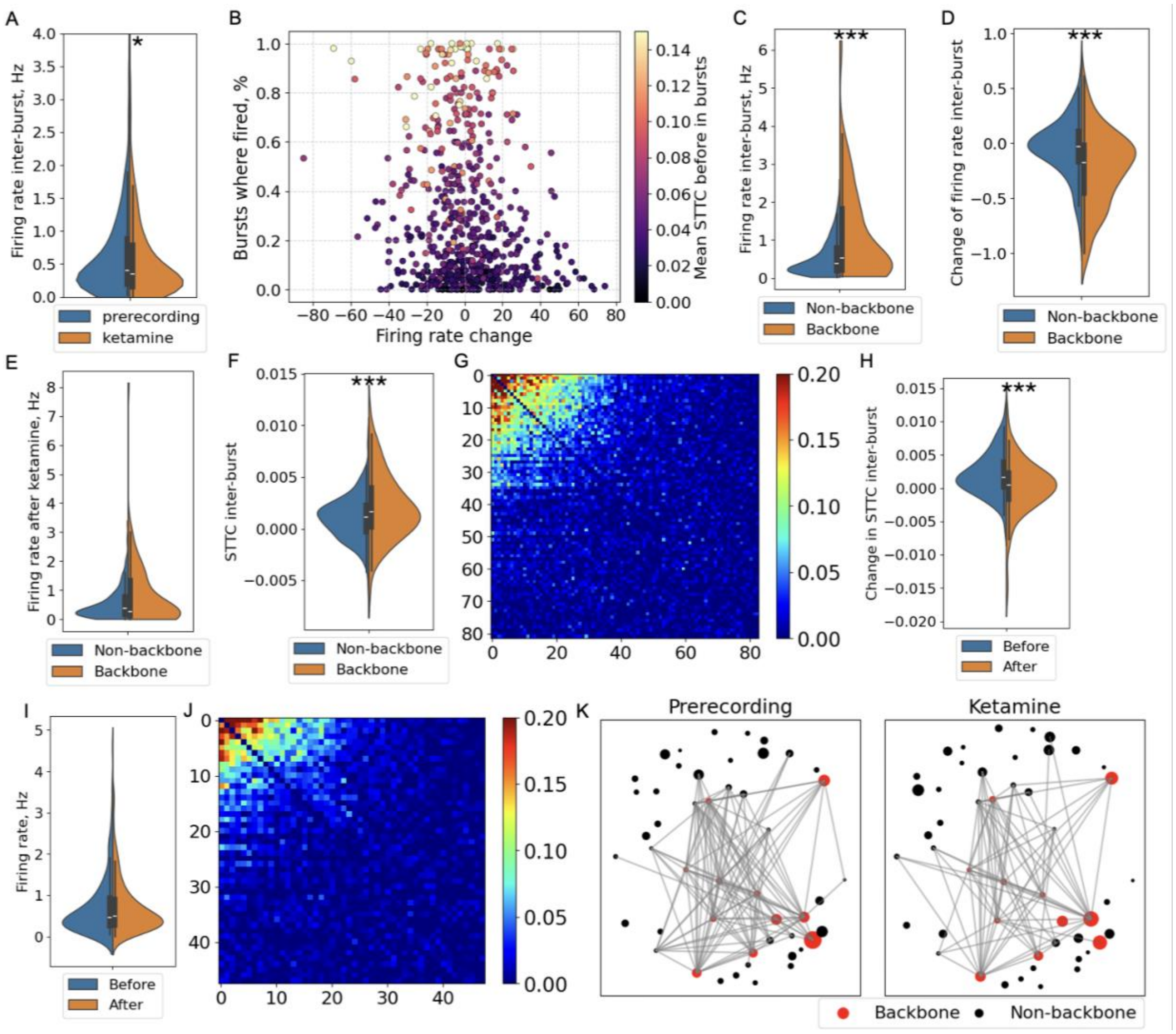


**Supplemental figure 2**

A. The distribution of unit firing rates before and after ketamine treatment in the inter-burst regions. B. While some firing rate fluctuations occur in saline-treated samples, there is no preferential suppression of the backbone units. C. Backbone units have higher firing rates than non-backbone units in the inter-burst regions. D. Backbone units are preferentially suppressed even in the inter-burst regions showed by bigger relative reduction of the firing rate. E. Firing persist in backbone units after ketamine treatment. F. Backbone units exhibit slightly but significantly higher STTC than non-backbone in the inter-burst regions. G. No reduction is STTC occurs after saline treatment. H. Backbone units show slight but significant reduction in STTC after ketamine treatment in the inter-burst regions. I. Firing rates are not suppressed by ketamine addition on day 7 of the chronic protocol. J. STTC values are not decreased by ketamine addition on day 7 of the chronic protocol. K. Functional connectivity graph doesn’t change with ketamine addition on day 7 of the chronic protocol.

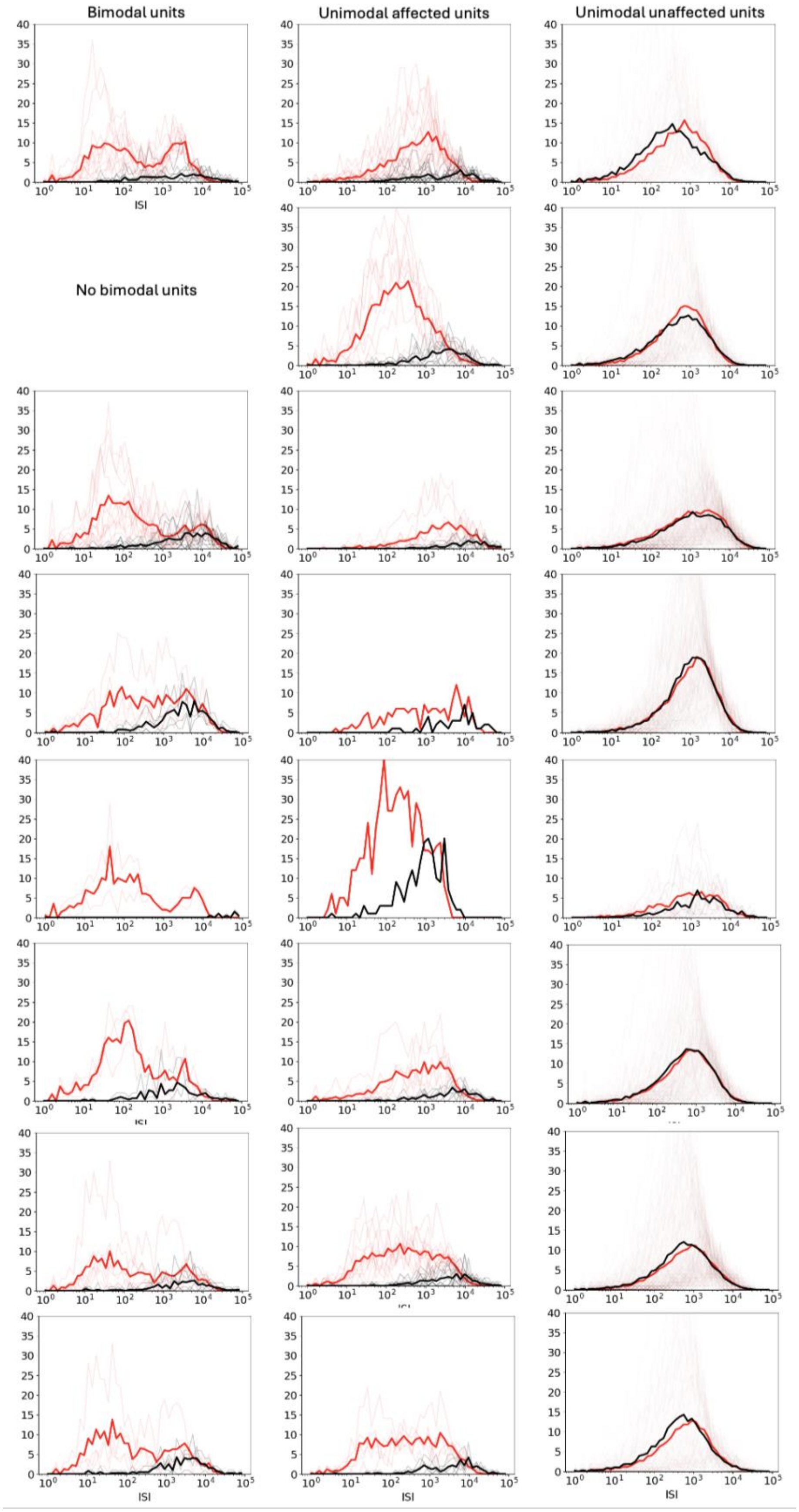


**Supplemental figure 3**. ISI distributions of bimodal and unimodal units. Each row represents an organoid.

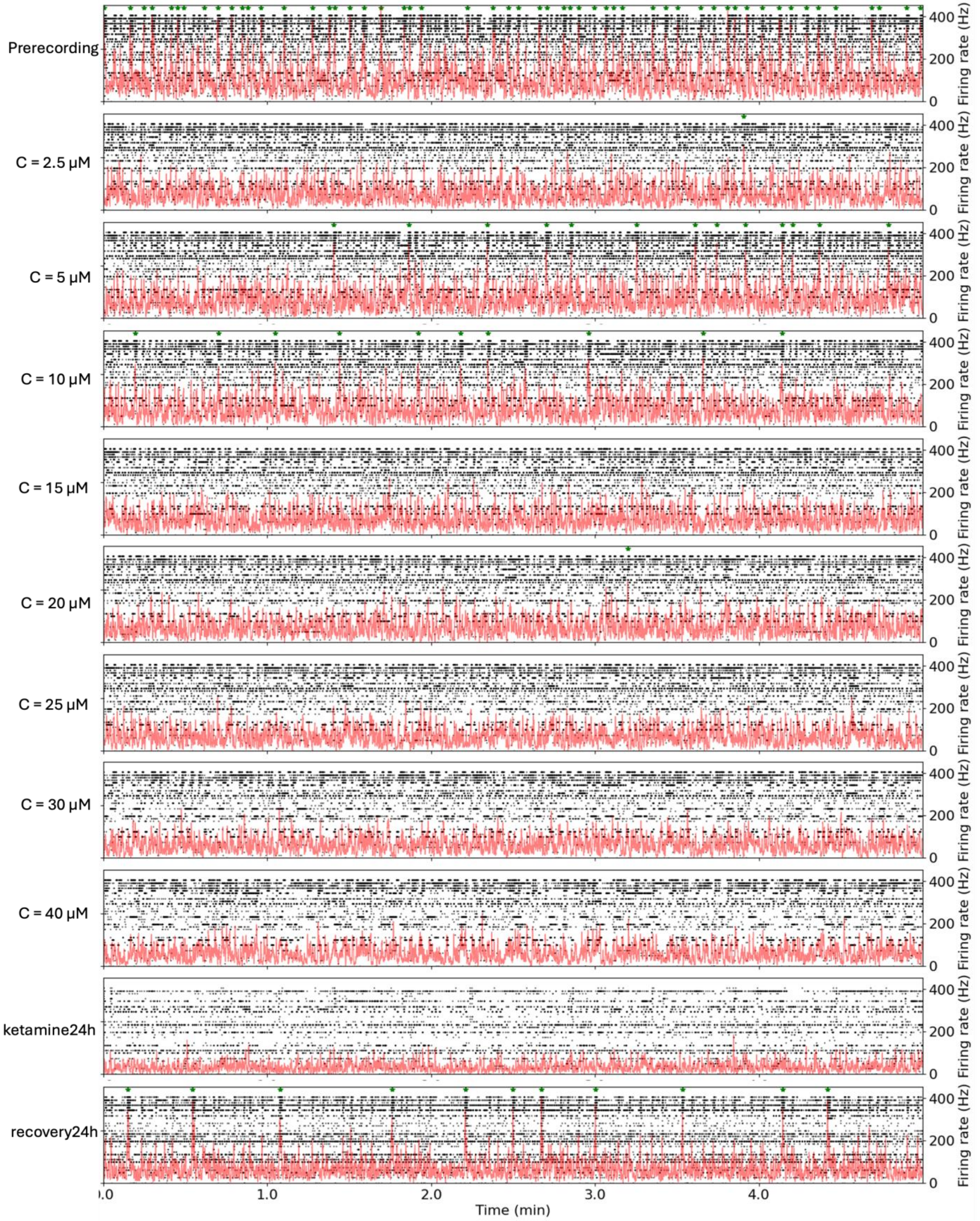


**Supplemental figure 4. Raster plots of ketamine dose responses from organoid 25497.**

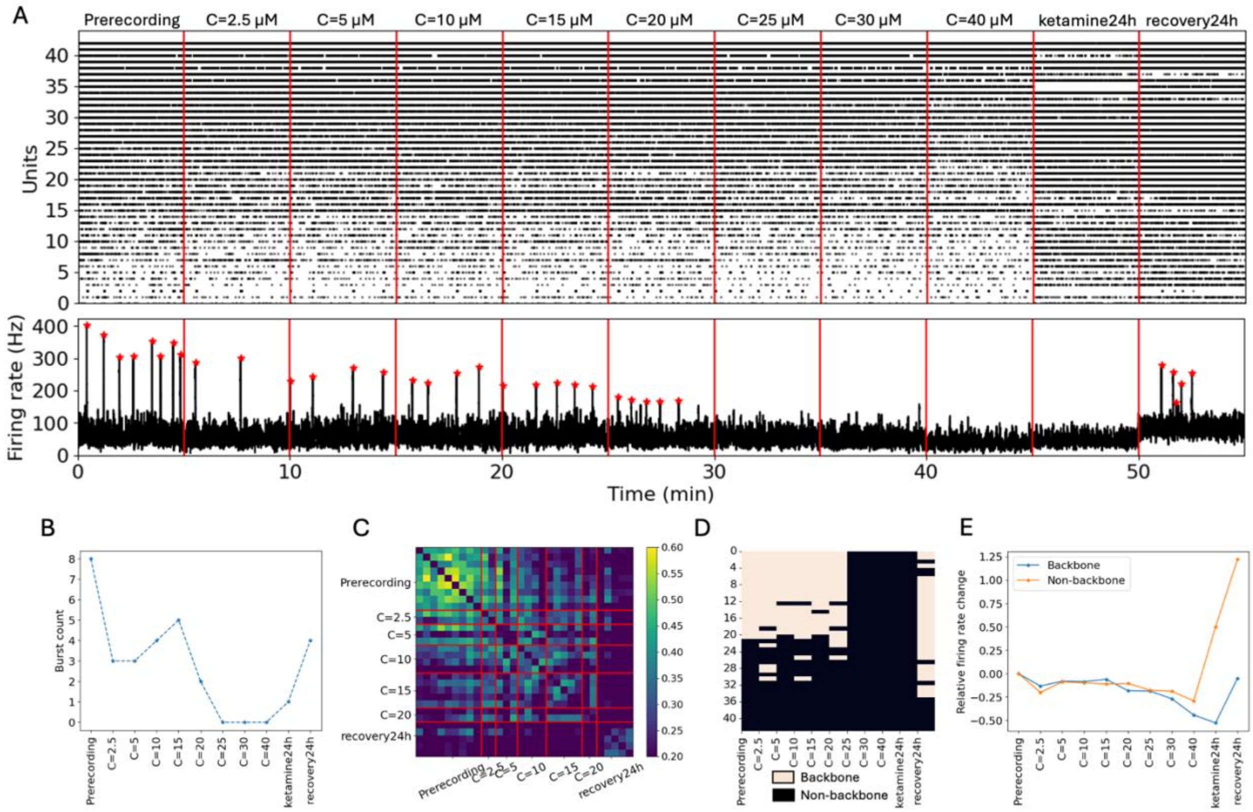


**Supplemental Figure 5. Ketamine dose–response and recovery, organoid 25509.** (A) Raster plot from a single organoid during eight consecutive ketamine dose steps (0–40 μM), followed by 24 h of continuous high-dose exposure and a subsequent 24 h recovery period in saline. Refer to Supp. Figure 6 for individual raster plots. (B) Total burst count as a function of dose. Bursting drops sharply after the first ketamine addition (2.5 μM), partially rebounds at intermediate doses, and returns after 24 h recovery in saline. (C) Burst-to-burst firing-rate cross-correlation indicates that burst structure does not return to its pre-exposure pattern after 24 h recovery. (D) Backbone units identified during the baseline recording largely remain backbone units throughout the dose–response protocol; additional backbone units emerge with increasing dose and after recovery. (E) Relative changes in mean firing rate show that backbone units reduce firing more rapidly than non-backbone units as dose increases.

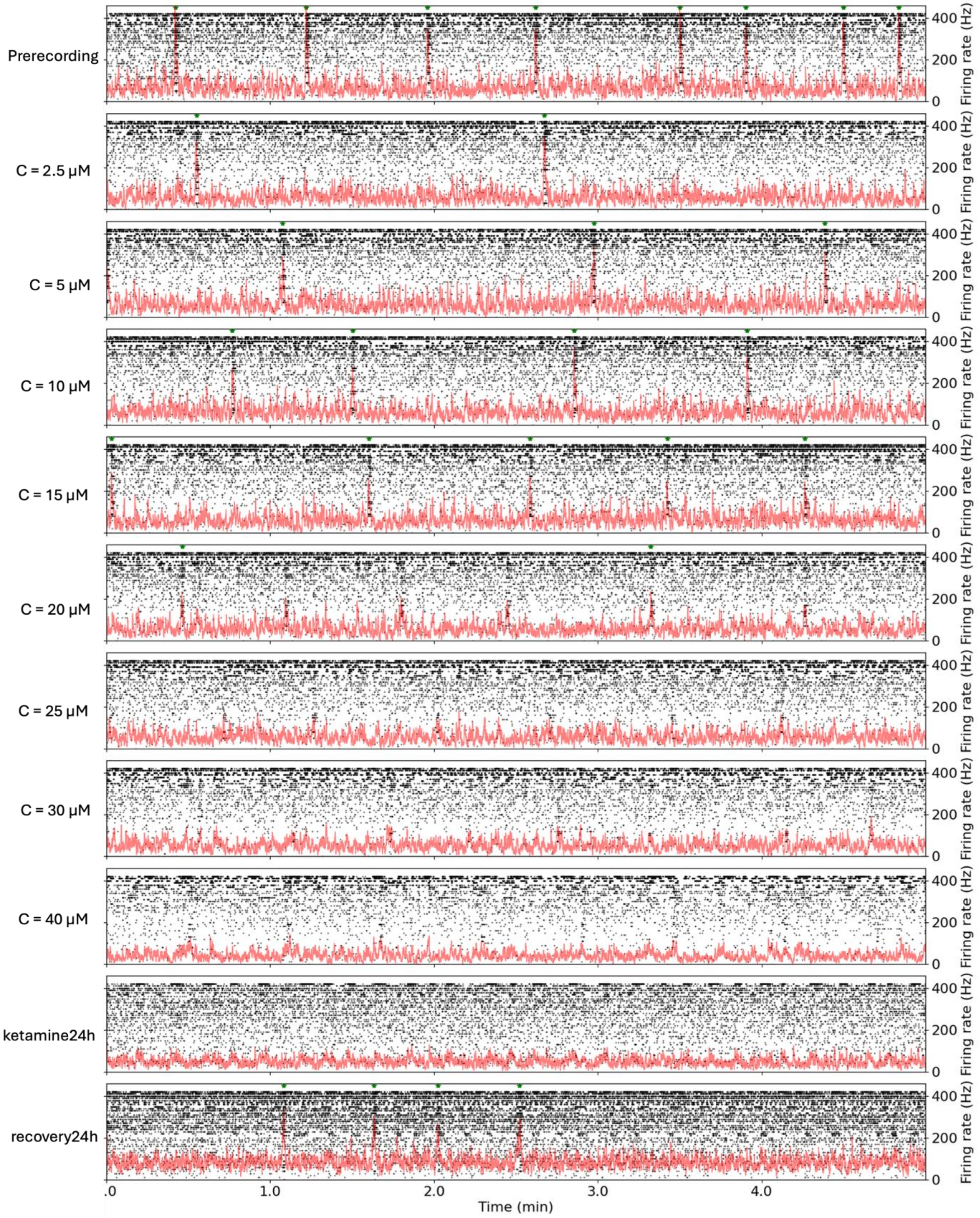


**Supplemental figure 6. Raster plots of ketamine dose responses from organoid 25509.**
